# From the clinic to the bench and back again in one dog year: identifying new treatments for sarcoma using a cross-species personalized medicine pipeline

**DOI:** 10.1101/517086

**Authors:** Sneha Rao, Jason A. Somarelli, Erdem Altunel, Laura E. Selmic, Mark Byrum, Maya U. Sheth, Serene Cheng, Kathryn E. Ware, So Young Kim, Joseph A. Prinz, Nicolas Devos, David L. Corcoran, Arthur Moseley, Erik Soderblom, S. David Hsu, William C. Eward

**Affiliations:** Department of Medicine, Duke University Medical Center, Durham, NC, USA; Duke Cancer Institute, Durham, NC, USA; Department of Veterinary Clinical Sciences, College of Veterinary Medicine, The Ohio State University, Columbus, OH, USA; Pratt School of Engineering, Duke University; Department of Molecular Genetics and Microbiology; Duke Center for Genomic and Computational Biology; Department of Orthopaedic Surgery

**Keywords:** precision medicine, cancer therapy, leiomyosarcoma, drug discovery, comparative oncology

## Abstract

Cancer drug discovery is an inefficient process, with more than 90% of newly-discovered therapies failing to gain regulatory approval. Patient-derived models of cancer offer a promising new approach to identifying personalized treatments; however, for rare cancers, such as sarcomas, access to patient samples can be extremely limited, which precludes development of patient-derived models. To address the limited access to patient samples, we have turned to pet dogs with naturally-occurring sarcomas. Although sarcomas make up less than 1% of all cancers in humans, sarcomas represent at least 15% of all cancers in dogs. Dogs with naturally-occurring sarcomas also have intact immune systems, an accelerated pace of cancer progression, and share the same environment as humans, making them ideal models that bridge key gaps between mouse models and human sarcomas.

Here, we develop a framework for a personalized medicine pipeline that integrates drug screening, validation, and genomics to identify new therapies. We tested this paradigm through the study of a pet dog, Teddy, who presented with six synchronous leiomyosarcomas. By integrating patient-derived cancer models, in vitro drug screens, and in vivo validation we identified proteasome inhibitors as a potential therapy for Teddy. After showing an initial response to the proteasome inhibitor, bortezomib, Teddy developed rapid resistance, and tumor growth resumed. Whole exome sequencing revealed substantial genetic heterogeneity across Teddy’s multiple recurrent tumors and metastases, suggesting that intra-patient heterogeneity was responsible for the heterogeneous clinical response. Ubiquitin proteomics coupled with exome sequencing revealed multiple candidate driver mutations in proteins related to the proteasome pathway. Together, our results demonstrate how the comparative study of canine sarcomas can offer rapid insights into the process of developing personalized medicine approaches that can lead to new treatments for sarcomas in both humans and canines.

## Introduction

Despite spending billions of dollars on the preclinical development of new anti-cancer drugs, fewer than 1 in 10 new therapies make it from the bench to the bedside and gain FDA approval (*1*). These sobering statistics clearly demonstrate that the preclinical models and paradigms currently being used to discover new cancer treatments require improvement. This need for improvement is exemplified by the slow progress in finding new therapies for sarcoma. Sarcomas are rare, but highly aggressive cancers that are prevalent in children and young adults. While sarcomas make up less than 1% of adult solid tumors, they account for nearly 15% of pediatric solid tumors(*2*). For patients who present with metastatic disease, the 5-year survival is just 16% (*3*). Few new therapies have emerged in recent decades, underscoring the need for creative new approaches in drug discovery.

One approach that has increasingly become a part of the discovery pipeline is the use of patient-derived models of cancer, including low-passage cell lines and patient-derived xenografts (PDXs). To create these patient-derived models, individual patient tumors are grown directly in culture or in immunocompromised mice. Each type of patient-derived model has unique advantages: For example, patient-derived cell lines enable large-scale drug screens to take place quickly and at low cost. On the other hand, the use of PDXs reduces the selective bottleneck of cell line generation and maintains the stromal components of the original tumor, which are increasingly recognized as critical components of a tumor’s relative therapeutic sensitivity(*4, 5*). These patient-derived models are also being used to develop personalized treatments and guide development of novel targeted agents (*6, 7*). One study in colorectal cancers showed a correlation between transplanted xenograft tumors and clinical response to cytotoxic therapy (*8*). Another pilot clinical trial of patients with advanced solid tumors received systemic cytotoxic therapies based on *in vivo* validation in PDXs (*9*). This study showed that 11 out of 17 treatment regimens identified in PDX were clinically efficacious (*10*). Drug screening in this study was done *in vivo* rather than *in vitro* and used over 200 treatment regimens, including both targeted and non-targeted agents (*10*). A similar study in advanced sarcoma patients with a variety of histologic subtypes also yielded concordant results between PDX and patient responses, with 13 out of 16 patients showing a correlation between efficacy of the top drug identified through PDX drug trials and clinical outcomes (*11*). Yet despite these exciting results, there remains a disconnect between drug testing in mice and performance in human patients.

Another approach for cancer drug discovery that is rapidly gaining attention is the study of pet dogs with spontaneously-occurring sarcomas and the inclusion of these patients in therapeutic trials. Canine sarcomas are far more prevalent than their human counterparts, representing approximately 15% of all canine malignancies (*12*) and rendering them an underutilized “model” of human disease (*13, 14*). Unlike mouse models – which often fail to recapitulate key conditions of spontaneous human disease - dogs share an environment with humans, have an intact immune system, and have nearly identical treatment options. While there are some differences in the histopathologic grading of soft tissue sarcomas between humans and dogs, a study using canine soft tissue sarcomas to compare pathologic diagnoses between veterinary and medical pathologists showed that the majority of canine tumors were given diagnoses congruent with the human counterpart (*15*). Coupled with patient-derived models and precision medicine strategies, a cross-species approach could illuminate new therapeutic options for sarcoma patients with greater fidelity than the traditional “cells, then mice, then humans” pathway. Most importantly, because the lifespan of dogs is much shorter than that of humans, discoveries in canine clinical trials can be made more quickly in canine patients given the rapid progression of their lives relative to humans.

In the present work, we report the development and testing of a personalized medicine pipeline that combines patient-derived models, personalized genomics, and drug screening strategies to identify new potential therapies for a young dog who presented with seven synchronous, spontaneous leiomyosarcomas. Using this pipeline, we first developed an early passaged cell line and PDX for our patient. Using high throughput drug screen on the cell line, we identified proteasome inhibitors as a candidate therapy for this patient, then validated the tumor response to proteasome inhibition *in vivo* using the patient’s PDX, and finally treated the patient’s recurrent tumor in the clinic with the proteasome inhibitor, bortezomib. Our work provides a generalizable framework for personalized medicine strategies and highlights key challenges in the development of such approaches.

## Materials and Methods

### Generation of patient-derived xenograft models

Tumor samples were collected from a three-year-old male golden retriever following surgical resection of the tumors at University of Illinois at Urbana-Champaign, College of Veterinary Medicine (Urbana, IL, USA) with the informed consent of the owner. PDX models of the patient’s sarcoma were generated as described previously, and all *in vivo* mouse experiments were performed in accordance with the animal guidelines and with the approval of the Institutional Animal Care and Use committee (IACUC) at the Duke University Medical Center (*16*). To develop PDXs, the tumor sample was washed in phosphate buffered saline (PBS), dissected into small pieces (<2 mm), and injected into the flanks of 8-10-week-old JAX NOD.CB17-PrkdcSCID-J mice obtained from the Duke University Rodent Genetic and Breeding Core. Tumors were passaged into successive mice once the tumor size reached between 500 to 1,500 mm^3^. Resected PDX tumors were homogenized in a PBS suspension and 150 µl of PDX tissue-PBS suspensions at 150 mg/ml concentration were injected subcutaneously into the right flanks of the 8 weeks old JAX NOD.CB17-PrkdcSCID-J mice. To maintain integrity of the PDX tumor, passages were limited to the 3rd generation.

### Low-passage cell line generation and characterization

Low passage cell lines were generated from the patient’s PDX during passage one of the PDX as follows. PDX tumor was surgically removed with a sterile blade, washed in PBS, and small pieces (< 2mm) of tumor tissue were mechanically homogenized and then suspended in cell growth media and cultured in 12-well plates with DMEM + 10% FBS + 1% Penicillin/Streptomycin. To isolate tumor cells, growing colonies of cells were isolated by trypsinization using O rings and cultured in fresh 12-well plates. This process was repeated until a colony of cells was established that resembled pure tumor cells in morphology. Contamination of the PDX cell line with mouse fibroblasts was detected by polymerase chain reaction (PCR) using canine-specific and mouse-specific primers. The following primers were used: canine reverse (5’-GTA AAG GCT GCC TGA GGA TAA G-3’), canine forward (5’-GGT CCA GGG AAG ATC AGA AAT G-3’), mouse reverse (5’-AGG TGT CAC CAG GAC AAA TG-3’), and mouse forward (5’-CTG CTT CGA GCC ATA GAA CTA A-3’) (*17*).

### High-throughput drug screening

Canine leiomyosarcoma low-passage cell line was cultured in DMEM + 10% FBS + 1% Penicillin/Streptomycin. Automated systems were used for a 119- and 2,100-compound high-throughput drug screens. The 119-drug screen library (Approved Oncology Set VI) was provided by the NCI Developmental Therapeutics Program (https://dtp.cancer.gov/). Automated liquid handling was provided by the Echo Acoustic Dispenser (Labcyte) for drug addition or Well mate (Thermo Fisher) for cell plating, and asays were performed using a Clarioscan plate reader (BMG Labtech). The BioActive compound library includes 2,100 small molecules that are annotated for pathway and drug target (Selleckchem) and was screened in triplicate. Compounds were stamped into 384 well plates for a final concentration of 1 μM using an Echo Acoustic Dispenser (Labcyte). Cells were then plated at a density of 2,000 cells/well using a WellMate (ThermoFisher) and incubated in the presence of drug for 72 hours. After 72 hours of incubation, Cell Titer Glo was added to each well and luminescence was measured using a Clariostar Plate Reader (BMG Labtech). Percent killing was quantified using the formula 100*(1-(average CellTiterGlo^drug^/average CellTiterGlo^DMSO^)) where the value average CellTiterGlo^DMSO^ was the average DMSO CellTiterGlo value across each plate.

### Validation of top drug candidates *in vivo*

To validate top candidates from the *in vitro* drug screens 150 μl of homogenized PDX tissue-PBS suspensions were injected at a concentration of 150 mg/ml of tumor tissue subcutaneously into the right flanks of the 8-10 weeks old JAX NOD.CB17-PrkdcSCID-J mice. Top drug targets identified by the high-throughput drug screens for *in vivo* validation, bortezomib (PS-341) and 17-DMAG (alvespimycin) HCl were purchased from Selleck Chemicals (Houston, TX). Drug were first solubilized in DMSO and then diluted in PBS for intraperitoneal injections. When the tumor volumes reached 100-150 mm^3^, mice were randomized (n = 5 mice for each treatment group) and 1 mg/kg bortezomib and 25 mg/kg alvespimycin intraperitoneal injections were initiated three times a week (*18, 19*). Control tumors were treated with 100*µ*l of 5% DMSO diluted in PBS. Tumor volumes were measured three times a week using calipers, and (length x (width)^2^)/2 was used to calculate the tumor size. Mice were sacrificed on day 18 or if the tumor volume reached 1,500 mm^3^.

### Whole exome sequencing

Genomic DNA from seven primary tumors, one recurrent tumor, a patient-derived xenograft, and the cell line were isolated using the QIAGEN DNeasy Blood and Tissue kit. DNA quality analysis, exome capture, and sequencing were performed at the Duke University Sequencing and Genomics Technologies Shared Resource. Genomic DNA samples were quantified using fluorometric quantitation on the Qubit 2.0 (ThermoFisher Scientific). For each sample, 1ug of DNA was sheared using a Covaris to generate DNA fragments of about 300bp in length. Sequencing libraries were prepared using the Roche Kapa HyperPrep Library prep Kit. During adapter ligation, unique indexes were added to each sample. Resulting libraries were cleaned using SPRI beads and quantified on the Qubit 2.0. Size distributions were checked on an Agilent Bioanalyzer. Libraries were pooled into equimolar concentration (8 libraries per pool) and library pools were finally enriched using the Roche SeqCap^®^ EZ Dog Exome panel (design 1000003560). Each pool of enriched libraries was sequenced on one lane of a HiSeq 4000 flow cell at 150bp PE, generating about 41 Million clusters per sample or ∼12Gb of data. Sequence data was demultiplexed and Fastq files generated using Bcl2Fastq2 conversion software provided by Illumina.

Initial data analysis and variant calling were performed by the Duke University Genomic Analysis and Bioinformatics Resource. Exome sequencing data was processed using the TrimGalore toolkit (*20*), which employs Cutadapt(*21*) to trim low-quality bases and Illumina sequencing adapters from the 3’ end of the reads. Reads were aligned to the CanFam3.1 version of the dog genome with the BWA algorithm (*22, 23*). PCR duplicates were flagged using the PICARD Tools software suite (*24*). Alignment processing and variant calling were performed using the MuTect2 (*25*) algorithm that is part of the GATK(*22*) following the Broad Institute’s Best Practices Workflow for identifying somatic variants(*22*). Variants for each sample were called relative to the normal sample. Variant call files for each sample were filtered for single nucleotide polymorphisms using the Genome Analysis Toolkit and converted to PHYLIP format using the vcf2phylip package (*27*). Phylogenetic trees were generated using PHYLIP with 1,000 bootstrap replicates per tree (*28*) and visualized using the ape package in R (*29*). The number of shared mutations was calculated pairwise between the matched tumor-normal variants of each sample using VCFtools (*30*). Genes with deleterious mutations in each sample were identified using Ensembl’s Variant Effect Predictor tool (*30*). These results were analyzed and visualized using BioVenn and the UpSetR package in R (*31, 32*).

### Ubiquitin-tagged proteomics analysis of PDX tumors treated with bortezomib

#### Sample Preparation

Flash frozen vehicle- and bortezomib-treated PDX tumors (n = 3 per treatment) were provided to The Duke Proteomics and Metabolomics Shared Resource for processing and analysis. Samples were normalized to 3.3 μL of 8 M urea per mg of wet weight and homogenized using a bead beater at 10,000 rpm. Protein concentration was determined via Bradford assay and was normalized to 5,000 μg of protein in 1.6 M of urea using 50 mM ammonium bicarbonate. Samples were then reduced with 10 mM dithiothreitol for 45 minutes at 32°C and alkylated with 25 mM iodoacetamide for 45 minutes at room temperature. Trypsin was added to a 1:25 ratio (enzyme to total protein) and allowed to proceed for 18 hours at 37°C. After digestion, peptides were acidified to pH 2.5 with trifluoroacetic acid (TFA) and subjected to C18 SPE cleanup (Sep-Pak, 50 mg bed).

For ubiquitin antibody enrichment, samples were resuspended in 750 uL 1X IAP Buffer (50 mM MOPS pH 7.2, 10 mM sodium phosphate, 50 mM NaCl from Cell Signaling Technology) using vortex and brief bath sonication. Pre-aliquoted PTMScan^®^ Pilot Ubiquitin Remant Motif (K-□-GG) beads (Cell Signaling Technology) were thawed for each sample, storage buffer was removed following slow centrifugation, and beads were pre-washed with 4 x 1 mL of 1X PBS buffer. Resuspended peptides were then transferred in IAP buffer directly onto beads. Immunoprecipitation was performed for 2 hours at 4C using end-over-end mixing. After spinning gently to settle the beads (VWR microfuge) the supernatants were removed. The IAP resins containing the enriched ubiquitinated peptides were then washed with 1mL of IAP buffer three times, and one time with 0.1X IAP buffer. After removing the supernatants, the antibody-bound ubiquitinated peptides were eluted with a 50 μl aliquot of 0.15% TFA in water for approximately 10 minutes at room temperature, tapping gently on the bottom of the tube a few times during elution to ensure mixing. Beads were eluted a second time with 45 μL of 0.15% TFA in water and added to the first elution. Combined eluents were lyophilized to dryness. Samples were resuspended in 35 μL 0.1% formic acid for a final cleanup on a C18 Stage Tip. All samples were then lyophilized to dryness and resuspended in 12 μL 1%TFA/2% acetonitrile containing 12.5 fmol/μL yeast alcohol dehydrogenase. From each sample, 3 μL was removed to create a QC Pool sample that was run periodically throughout the acquisition period.

Quantitative LC/MS/MS was performed on 4 μL of each sample, using a nanoAcquity UPLC system (Waters Corp) coupled to a Thermo QExactive HF-X high resolution accurate mass tandem mass spectrometer (Thermo) via a nanoelectrospray ionization source. Briefly, the sample was first trapped on a Symmetry C18 20 mm × 180 μm trapping column (5 μl/minute at 99.9/0.1 v/v water/acetonitrile), after which the analytical separation was performed using a 1.8 μm Acquity HSS T3 C18 75 μm × 250 mm column (Waters Corp.) with a 90-minute linear gradient of 5 to 30% acetonitrile with 0.1% formic acid at a flow rate of 400 nanoliters/minute (nL/min) with a column temperature of 55°C. Data collection on the QExactive HF mass spectrometer was performed in a data-dependent acquisition (DDA) mode of acquisition with a r=120,000 (@ m/z 200) full MS scan from m/z 375 – 1600 with a target AGC value of 3e6 ions followed by 30 MS/MS scans at r=15,000 (@ m/z 200) at a target AGC value of 5×10^4^ ions and 45 ms. A 20 second dynamic exclusion was employed to increase depth of coverage. The total analysis cycle time for each sample injection was approximately 2 hours.

Data was imported into Proteome Discoverer 2.2 (Thermo Scientific Inc.), and analyses were aligned based on the accurate mass and retention time of detected ions using Minora Feature Detector algorithm in Proteome Discoverer. Relative peptide abundance was calculated based on area-under-the-curve of the selected ion chromatograms of the aligned features across all runs. The MS/MS data was searched against the TrEMBL *C. familiaris* database (downloaded in Nov 2017) with additional proteins, including yeast ADH1, bovine serum albumin, as well as an equal number of reversed-sequence “decoys”) false discovery rate determination. Mascot Distiller and Mascot Server (v 2.5, Matrix Sciences) were utilized to produce fragment ion spectra and to perform the database searches. Database search parameters included fixed modification on Cys (carbamidomethyl) and variable modifications on Lysine (Gly-Gly), and Meth (oxidation). Peptide Validator and Protein FDR Validator nodes in Proteome Discoverer were used to annotate the data at a maximum 1% protein false discovery rate.

### Data analysis and statistics

JMP from SAS software (Cary, NC, USA) was used for the high-throughput drug screen data analysis. Hierarchical clustering of data was used to identify the top drug candidates from the 119-compound drug screen and the 2,100-compound screen. Tumor volumes were recorded in GraphPad Prism 6 software (La Jolla, CA, USA). Two-way ANOVA analysis was used to compare differences in tumor volumes between the control and treatment groups.

## Results

### Applying a personalized medicine pipeline to an unusual case of leiomyosarcoma

We enrolled a three-year-old Golden Retriever (Teddy) for this study who presented to a veterinary primary care hospital with six synchronous leiomyosarcomas that underwent excisional biopsy (**Figure 1**). Teddy was then referred to the Small Animal Oncology team at the University of Illinois at Urbana-Champaign for treatment of a mass near the stifle. This tumor was excised and scars of the resected tumors were excised. During clipping and preparation for these surgeries, the treating surgeon noted two new masses in addition to previous surgical scars that were also resected and also determined to be high grade leiomyosarcoma (**Figure 1)**. Pathology reports from the time of tumor excision noted an “ulcerated, inflamed, highly cellular, invasive mass composed of neoplastic spindyloid cells arranged in short interlacing streams and bundles with many neutrophils throughout the neoplasm with clusters of lymphocytes and plasma cells at the periphery”, which was consistent with high grade leiomyosarcoma. Following surgery, Teddy was started on empirical treatment with toceranib, a multi-receptor tyrosine kinase inhibitor and the only FDA-approved targeted cancer therapeutic for dogs, given the high risk for recurrent disease.

**Figure 1.**
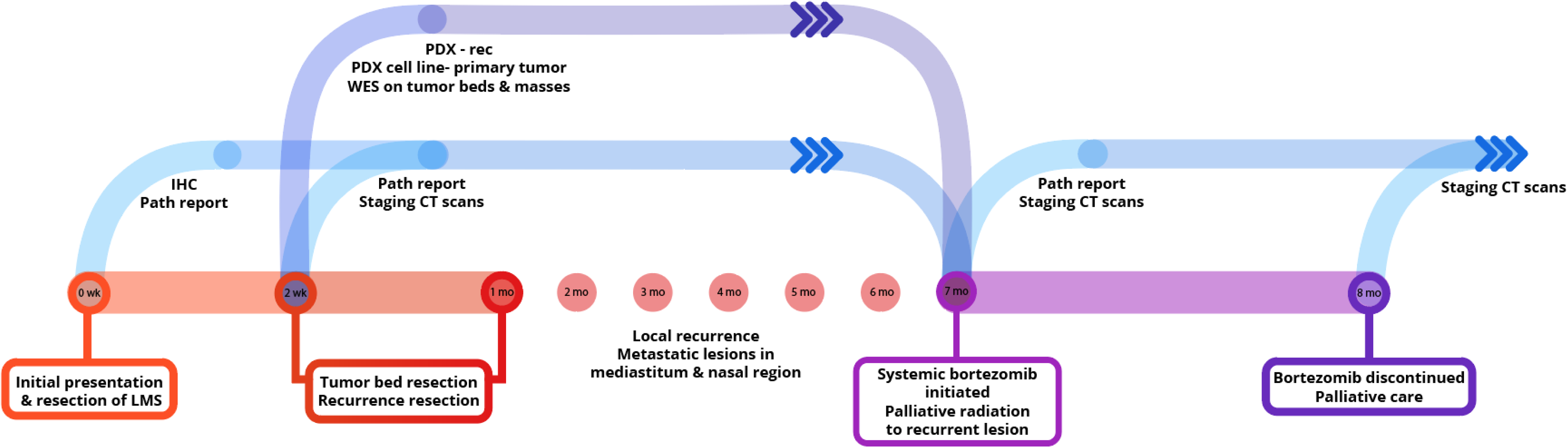
An integrated preclinical drug discovery and validation pipeline. A three year old canine patient with synchronous leiomyosarcomas (LMS) was identified and recruited based on high risk of disease recurrence. Using both *in vitro* and *in vivo* patient-derived models, we identified proteasome inhibitors as candidates for validation in clinic. Clinicians applied the information from this preclinical pipeline for the treatment of the patient’s recurrent and metastatic disease.

### Generation of patient-derived models of LMS-D48X

Using one of the excised recurrent tumors from this patient, we applied a personalized medicine pipeline to identify new potential therapies in the event that Teddy’s disease would eventually recur (**Figure 2A**). The pipeline included successful development of a matching PDX (designated “LMS-D48X”) and low-passage cell line, a high throughput drug screen on the cell line, genomic profiling of mutations in the original tumors, PDX, and cell line, and *in vivo* validation of top drug candidates (**Figure 2A**). Hematoxylin and eosin staining of the canine PDX revealed sheets of highly proliferative, spindle-like cells (**Figure 2B**). Similarly, the matched cell line was also highly proliferative, with an estimated doubling time of 26-36 hours and the presence of spindle-shaped, mesenchymal-like cells (**Figure 2C**). PCR using canine- and mouse-specific primers demonstrated that the LMS-D48X cell line is made up of purely canine tumor cells (**Figure 2D)**.

**Figure 2.**
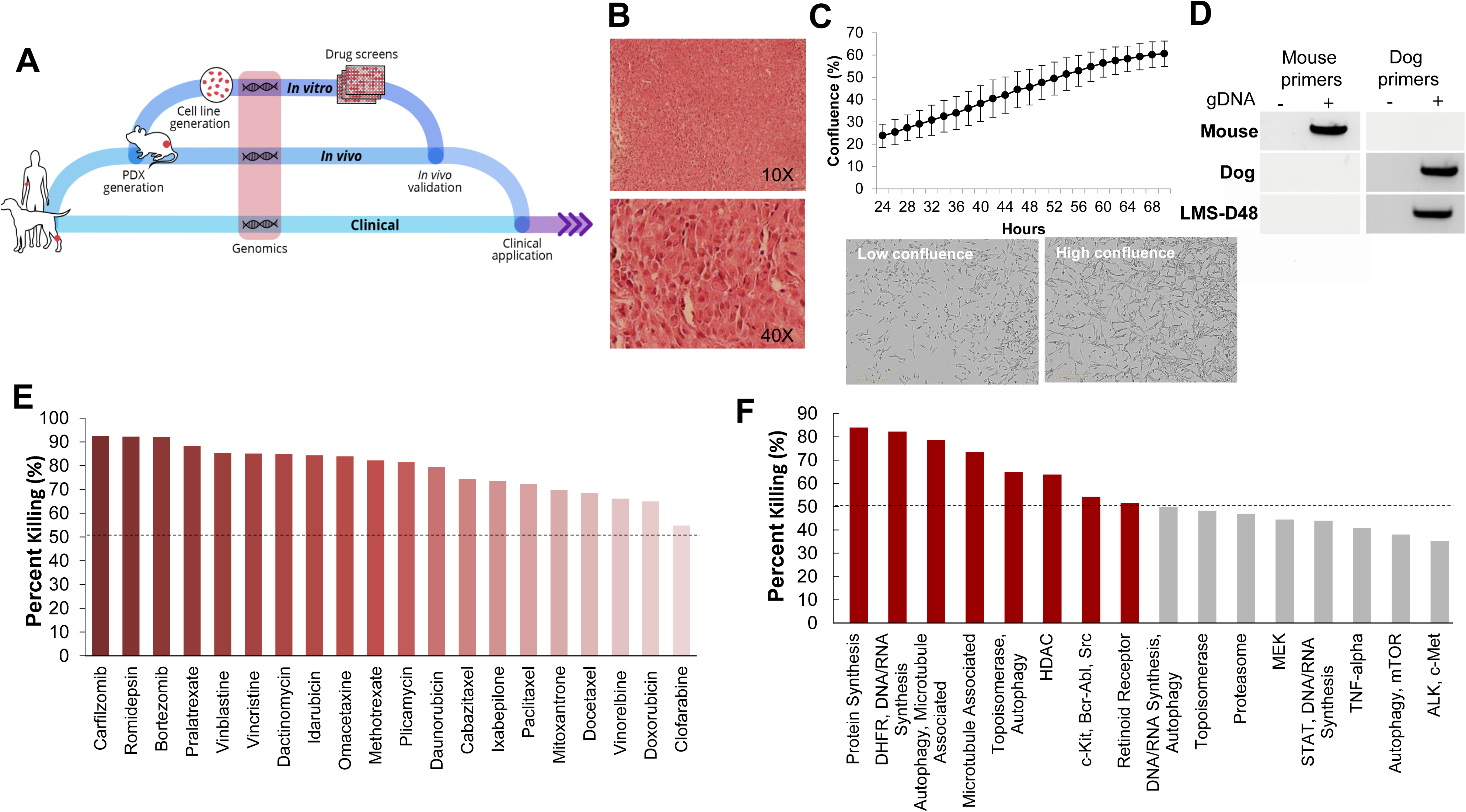
Patient-derived models of cancer enable seamless integration of high throughput drug screening with *in vivo* validations. **A.** Schematic of the personalized medicine pipeline integrating *in vitro* and *in vivo* drug discovery and validation. **B.** Hematoxylin and eosin stain of the patient derived xenograft model of the canine patient (LMS-D48) showing highly proliferative spindle-like cells. **C.** The patient-derived cell line also displays a high proliferation rate, with an estimated doubling time of 26-36 hours, and spindle-like mesenchymal morphology. **D.** A species-specific PCR using mouse- and canine-specific primers confirms that the patient-derived cell line is of canine origin. **E.** A preliminary drug screen of 119 FDA-approved compounds in the LMS-D48 cell line identified single standard-of-care agents and novel drug candidates. **F.** Analysis of drug screen data at the pathway level showed sensitivity to protein synthesis, DNA/RNA synthesis, autophagy, and HDAC inhibitors. Novel agents, including HDAC inhibitors and proteasome inhibitors were identified as top candidates for validation.

### High-throughput drug screens identify proteasome inhibitors as a potential candidate therapy

To identify potential candidate therapies to treat Teddy, we performed two high-throughput drug screens. First, we used a panel of 119 FDA-approved anti-cancer drugs. Importantly, this screen identified multiple standard-of-care therapies for soft tissue sarcomas, such as doxorubicin and danurubicin **(Figure 2E)**. Interestingly, however, in addition to standard-of-care therapies, the drug screen also identified several novel candidate drugs, such as proteasome inhibitors, HDAC inhibitors (i.e. romidepsin), and MEK inhibitors, as candidate agents (**Figure 2E**). Analysis of drug hits grouped by pathway revealed sensitivity to protein and nucleic acid synthesis pathways, autophagy, topoisomerases, HDACs, and c-kit/BCR/ABL (**Figure 2F**).

To further identify and validate additional novel therapeutic targets, we next performed a second-high throughput drug screen, this time using a larger panel of 2,100 bioactive compounds. The BioActives compound library (Selleckchem) contains a mixture of FDA-approved and non-FDA approved small molecules with confirmed bioactivity against known protein or pathway targets. The Bioactives collection is structurally diverse and is designed to target many key pathways regulating cellular processes including proliferation, apoptosis and signal transduction. Using the targeted pathway annotation for each compound, we were able to select targets and pathways for which multiple drugs had significant inhibitory effects. We hypothesized that this strategy would increase the likelihood of identifying the candidate targets/pathways for which a given tumor is most vulnerable. Our initial analysis of the screen revealed that a large portion (>90%) of compounds had little to no inhibitory effect, with only 6.6% of compounds showing >50% inhibition and 4.2% of drugs showing >75% inhibition **(Figure 3A)**. Analysis of top hits by cellular target demonstrated vulnerability for this cell line to some targets already identified from the 119-drug screen, such as proteasome inhibitors and MEK inhibitors, as well as novel drug classes, such as HSP, PLK, CRM1, NAMPT, Kinesin, and p53 inhibitors (**Figure 3B).** Analysis of the top inhibitors by pathway revealed enrichment in drugs targeting cytoskeletal signaling, the proteasome, apoptosis, cell cycle, and NF-κB (**Figure 3C**).

**Figure 3.**
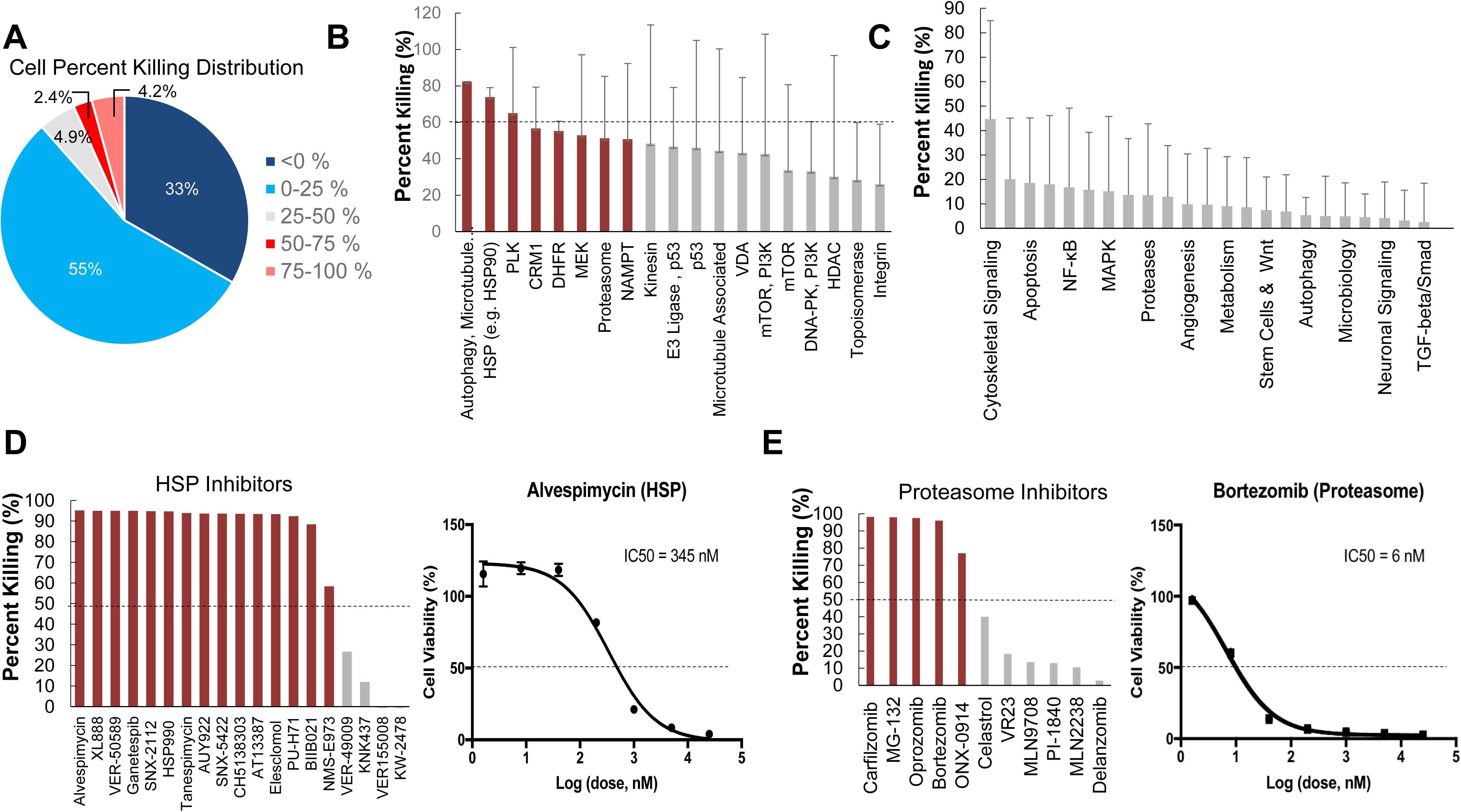
High-throughput drug screens identify HSP inhibitors and proteasome inhibitors as promising therapies for personalized treatment. **A.** LMS-D48 cells were plated at a density of 2,000 cells/well on plates prestamped with 2,100 drug compounds and DMSO. Cell titer glow assays were performed 72 hrs after cell plating to determine cell percent killing based on luminosity values. **B.** Analysis of drug targets from the 2,100 screen with multiple drugs shows HSP inhibitors and proteasome inhibitors among the top pathways for which this cell line displays significant sensitivity. **C.** Analysis of cellular pathways targeted by all drugs in the 2,100 drug screen shows that the cytoskeletal signaling pathway has the highest cell percent killing. **D.** LMS-D48 cells were sensitive to 15 out of 19 HSP inhibitors. Among these, alvespimycin was the top candidate, with an estimated IC_50_ of 345nM. **E.** Bortezomib was among the top drugs in the proteasome inhibitor class that killed LMS-D48 cells, with an estimated IC_50_ value of 6 nM.

We further explored the potential therapeutic efficacy of top pathways by analyzing the number of inhibitors for each pathway that had >50% cell growth inhibition. Notably, both the HSP and proteasome pathways had multiple drugs with >50% inhibition (15/19 and 5/11, respectively) (**Figure 3D and E**). In the proteasome inhibitor class, 4/11 drugs conferred >90% cell growth inhibition. Likewise, in the HSP inhibitor drug class, 13 out of 19 drugs caused >90 % cell growth inhibition **(Figure 3D and E)**. From these two drug classes, we selected alvespimycin (HSP inhibitor) and bortezomib (proteasome inhibitor) for further study. Both of these drugs have known toxicity profiles, with bortezomib being FDA approved for the treatment of multiple myeloma. *In vitro* validation of alvespimycin and bortezomib showed sub-micromolar IC_50_ values of 345 nM and 6nM, respectively (**Figure 3D and E)**.

### In vivo validation of alvespimycin and bortezomib in PDX models of LMS-D48X

We next used the LMS-D48X PDX to assess whether the top candidate therapies we identified *in vitro* would be therapeutically active in the patient’s matched PDX *in vivo*. Interestingly, while alvespimycin showed >95% growth inhibition *in vitro*, the PDX was unresponsive to this HSP inhibitor, with no difference in growth rate between vehicle-treated and alvespimycin-treated tumors (**Figure 4A**). On the other hand, tumors treated with bortezomib showed significant tumor growth inhibition, consistent with the *in vitro* drug screen (**Figure 4B, C).** Animal weights in LMS-D48 PDX mice did not change significantly from the vehicle-treated tumors in either of the drug treatment groups (**Figure 4D)**.

**Figure 4.**
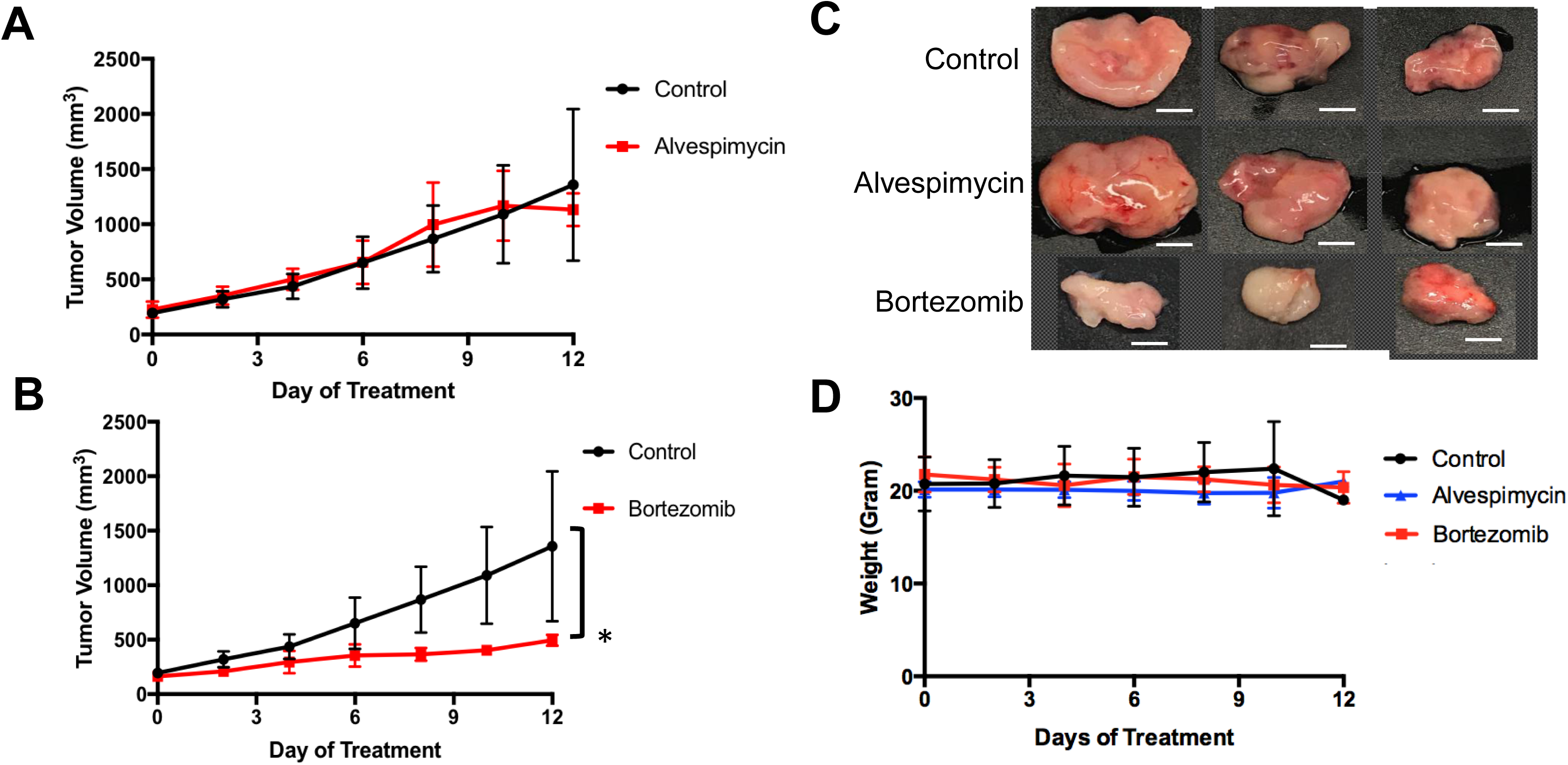
*In vivo* validation of top drug candidates reveals sensitivity of the LMS-D48 PDX to proteasome inhibition. **A.** Alvespimycin (25 mg/kg) was administered intraperitoneally (i.p.) *in vivo* to SCID beige mice harboring LMS-D48 PDX tumors (n = 5 mice per treatment group) each in control and treatment groups. There was no statistical difference between control and treatment groups as measured by analysis of variance. **B.** Bortezomib (1 mg/kg) was administered i.p. as described for alvespimycin above. Bortezomib significantly inhibited tumor growth of the PDX (p<0.0001). **C.** Representative images of resected tumors at treatment endpoint from the treatment and control groups show that control tumors are approximately twice the size of bortezomib-treated tumors (scale bar = 0.5 cm). **D.** Animal weights were not significantly changed during treatment with either alvespimycin or bortezomib during the treatment course.

### From bench to bedside: Applying preclinical modeling to clinical practice

For any personalized medicine approach to be clinically useful, it must provide insight into the patient’s disease within the time scale of clinical decision making. With an aggressive disease course and high likelihood for recurrence, Teddy presented a unique opportunity to assess the ability of our personalized medicine pipeline to meet the clinical demand for rapidly providing data on potential therapies to treating clinicians. Teddy presented at a six month follow up visit with lesions in the mediastinal and right iliac lymph nodes, nasal mucosa, and local recurrence in the right pelvic limb (**Figure 1 and Supplementary Figure 1)**. Using the *in vitro* screening and *in vivo* validations data from our pipeline, a decision was made to treat the patient with systemic bortezomib. The patient was treated with intravenous bortezomib infusions at 1.3mg/m^2^ twice weekly for four weeks and also received local palliative radiation therapy to the right pelvic limb to alleviate pain associated with the limb lesion. Measurements of the right pelvic limb lesion showed an initial decrease in tumor size during the first three weeks of treatment; however, tumor growth resumed by the sixth week of treatment (**Figure 5A)**. Metastatic lesions in other locations also increased in size on CT imaging at the conclusion of bortezomib treatment **(Figure 5B)**. Representative images of the tumors before and after bortezomib demonstrated the increase in tumor size and aggressive disease, especially in the infiltrative nature of the nasal mucosal lesion eroding into the maxilla (**Supplementary Figure 1)**.

**Figure 5.**
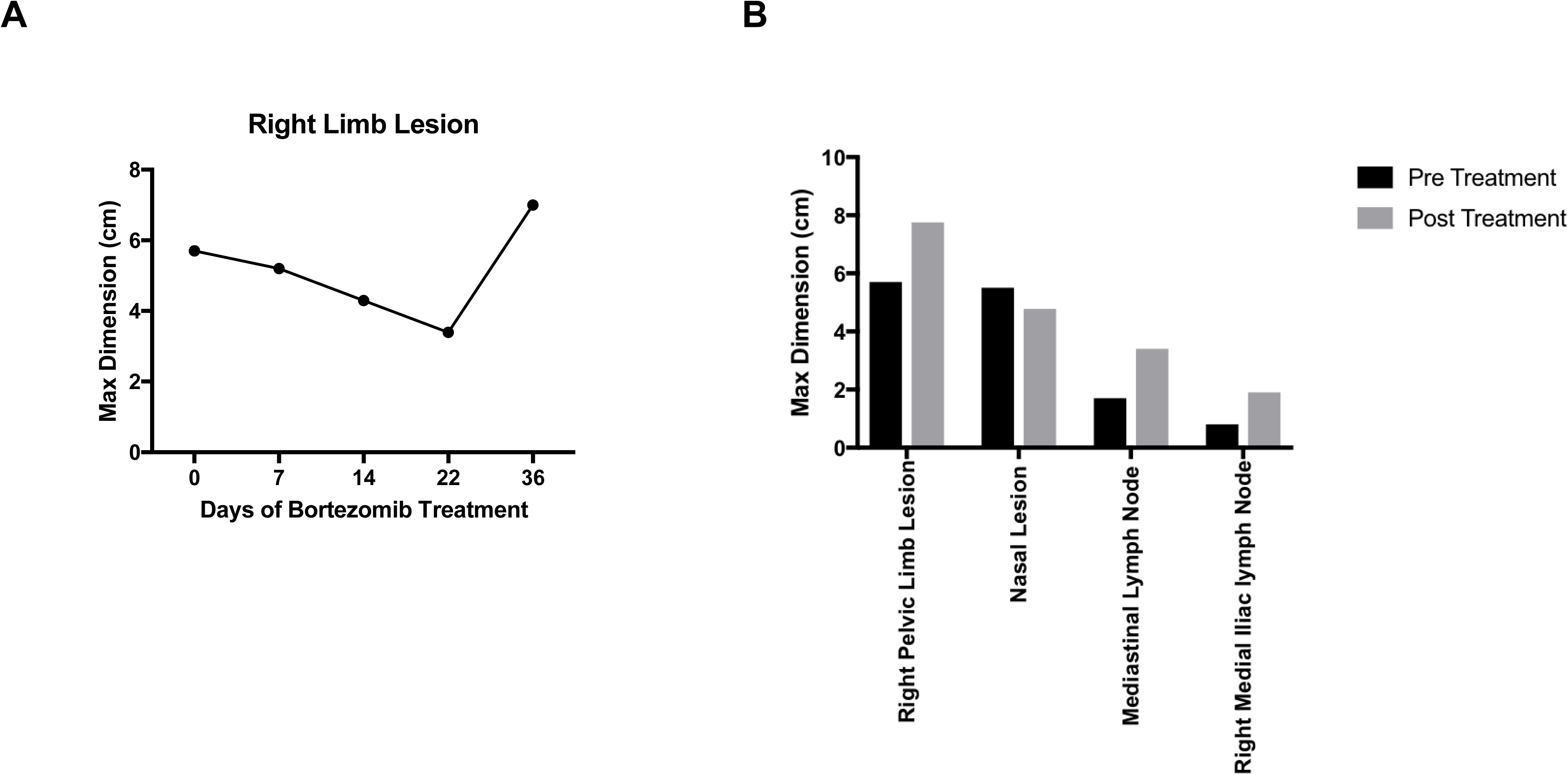
Translation of bortezomib into clinic. **A.** At the time of metastatic spread of disease, the patient had lesions in the mediastinal and right iliac lymph nodes, the nasal mucosa, and local recurrence at the right pelvic limb. The patient was started on systemic bortezomib therapy at a dose of 1.3mg/m2 twice weekly and palliative radiation therapy of 8 Gy by four fractions, once weekly, for pain from the right pelvic limb lesion. Measurement of the pelvic limb lesion during therapy showed decrease in maximal tumor dimension throughout 3 weeks of radiation therapy and systemic bortezomib; though, there was an increase in size two weeks after both therapies were stopped. **B.** CT staging studies and physical exam demonstrated an interval increase in tumor size at all sites of disease and after discontinuation of bortezomib therapy. The canine patient was then transitioned to palliative care.

### Whole exome sequencing reveals extensive inter-tumoral heterogeneity

Our analysis of patient-derived models of cancer identified bortezomib as a promising treatment for Teddy. Consistent with these preclinical observations, Teddy showed an initial response to bortezomib in the first three weeks of treatment. However, this response was short lived and tumor growth resumed however, Teddy also developed rapid resistance to systemic bortezomib by day 36 of treatment (**Figure 5A**). Given the substantial differences in response between tumor sites, we sought to better understand the underlying genetic landscape of the patient’s tumors and the relationship between these tumors and our patient-derived models. To do this, we performed whole exome sequencing and phylogenetic reconstructions on 11 samples from Teddy, including seven primary tumors, one recurrent tumor, one PDX and matched cell line, and normal tissue. Phylogenetic analysis of the tumors and patient-derived models grouped the PDX and cell line with the recurrent tumor with strong bootstrap support (**Figure 6A and Supplementary Figure 2**). With the exception of the distance trees, the grouping of the PDX and cell line with the recurrent tumor was consistent for all other methods of phylogenetic inference, including DNA compatibility, maximum parsimony, and maximum likelihood (**Figure 6B**). We also counted the number of shared somatic mutations across all samples and found the greatest similarity between the PDX, cell line, the recurrent tumor, and tumor 1 (**Figure 6C**). Together, these results suggest that the PDX and cell line most closely resemble the recurrent tumor. All other tumor samples shared little genetic overlap (3% to 16%). Tumor 7 was particularly distinct from the other tumors, sharing just 3.5% of somatic mutations with all other tumors (**Figure 6C**). Analysis of unique and shared somatic mutations revealed that unique mutations dominate the genetic landscape of each tumor (**Figure 6D**).

**Figure 6.**
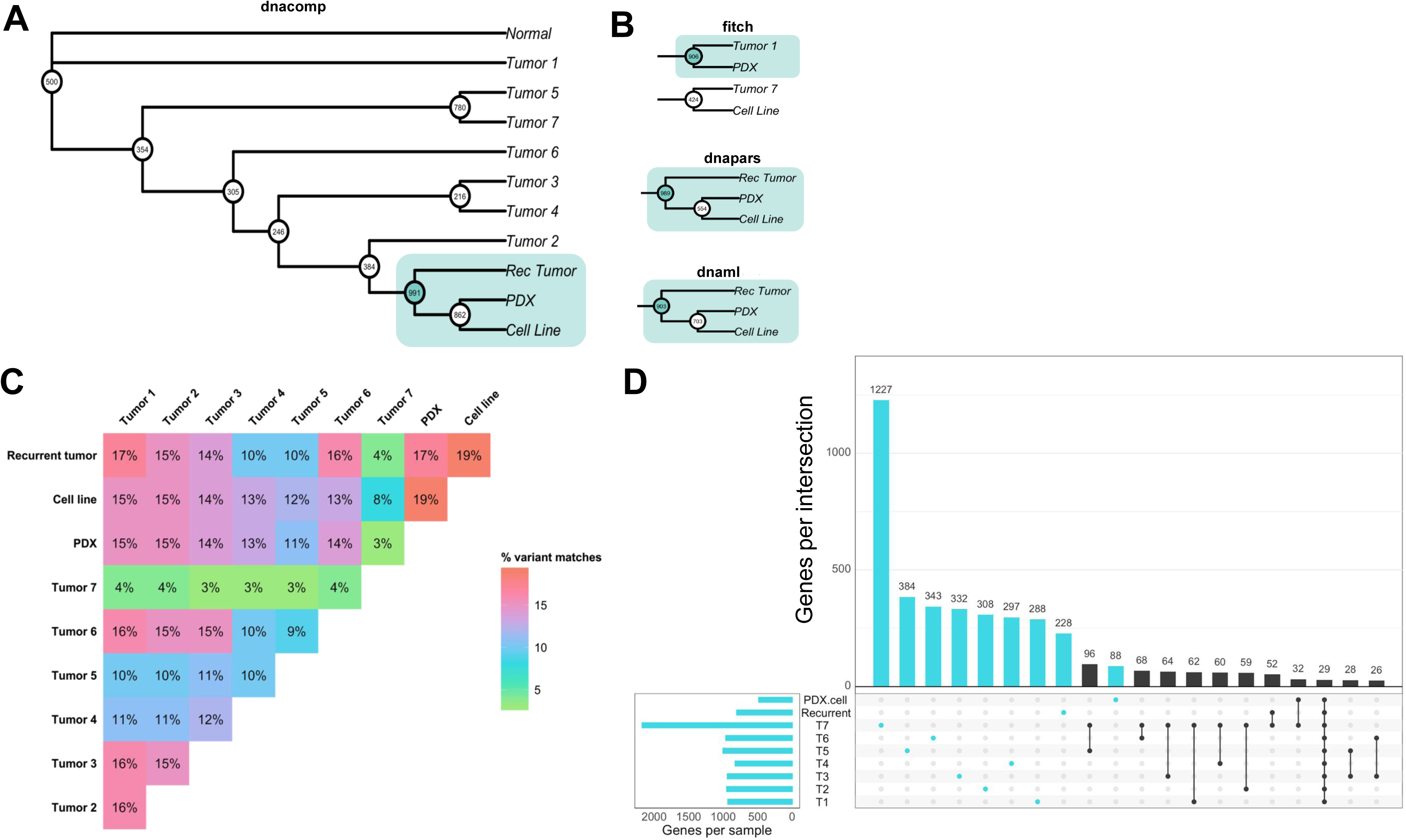
Whole exome sequencing reveals inter-tumoral heterogeneity across the patient’s tumors. **A.** Phylogenetic reconstruction using the DNA compatibility algorithm supports a clade that includes the PDX, cell line, and the recurrent tumor with bootstrap support greater than 900/1000. **B.** With the exception of the distance tree (Fitch), trees based on maximum parsimony and maximum likelihood also grouped the cancer models with the recurrent tumor. **C.** A similarity matrix comparing all somatic variants from each sample shows the percentage of shared mutations across all samples. **D.** Individual samples had higher numbers of mutated genes that were unique to each sample. Common shared mutations were relatively rare, reflecting the heterogeneity of the samples.

### Integration of whole exome sequencing and ubiquitin proteomics identifies potential mechanisms of action of bortezomib

To further understand the underlying molecular mechanisms of sensitivity and resistance to bortezomib for this patient, we performed mass spectrometry proteomics analysis of ubiquitin-tagged proteins in PDX tumors treated with vehicle or bortezomib. Since bortezomib is a proteasome inhibitor, we analyzed proteins that were differentially ubiquitinated in the PDX treated with bortezomib as compared to vehicle-treated tumors. We identified a total of 290 differentially ubiquitinated proteins in vehicle-vs. bortezomib-treated PDX tumors (adjusted p-value <0.05), 160 of which showed increased ubiquitination and 130 of which showed decreased ubiquitination (**Figure 7A**). Analysis of differentially ubiquitinated targets revealed enrichment for myosins and HSPs as the proteins with the greatest increase in ubiquitination in bortezomib-treated tumors as compared to vehicle-treated tumors (**Figure 7A and Supplementary File 1**). It is worth noting that the top hits were unique to this PDX, as additional proteomics analysis of bortezomib-treated osteosarcoma PDXs yielded a different suite of ubiquitinated proteins (Altunel et al., in preparation). Pathway analysis of proteins with increased ubiquitination revealed enrichments in pathways related to actin, contractile filament movement, and the proteasome (**Figure 7B**) and pathways related to proteins with decreased ubiquitination were enriched for adherens junctions, focal adhesions, and extracellular vesicles (**Figure 7C**).

**Figure 7.**
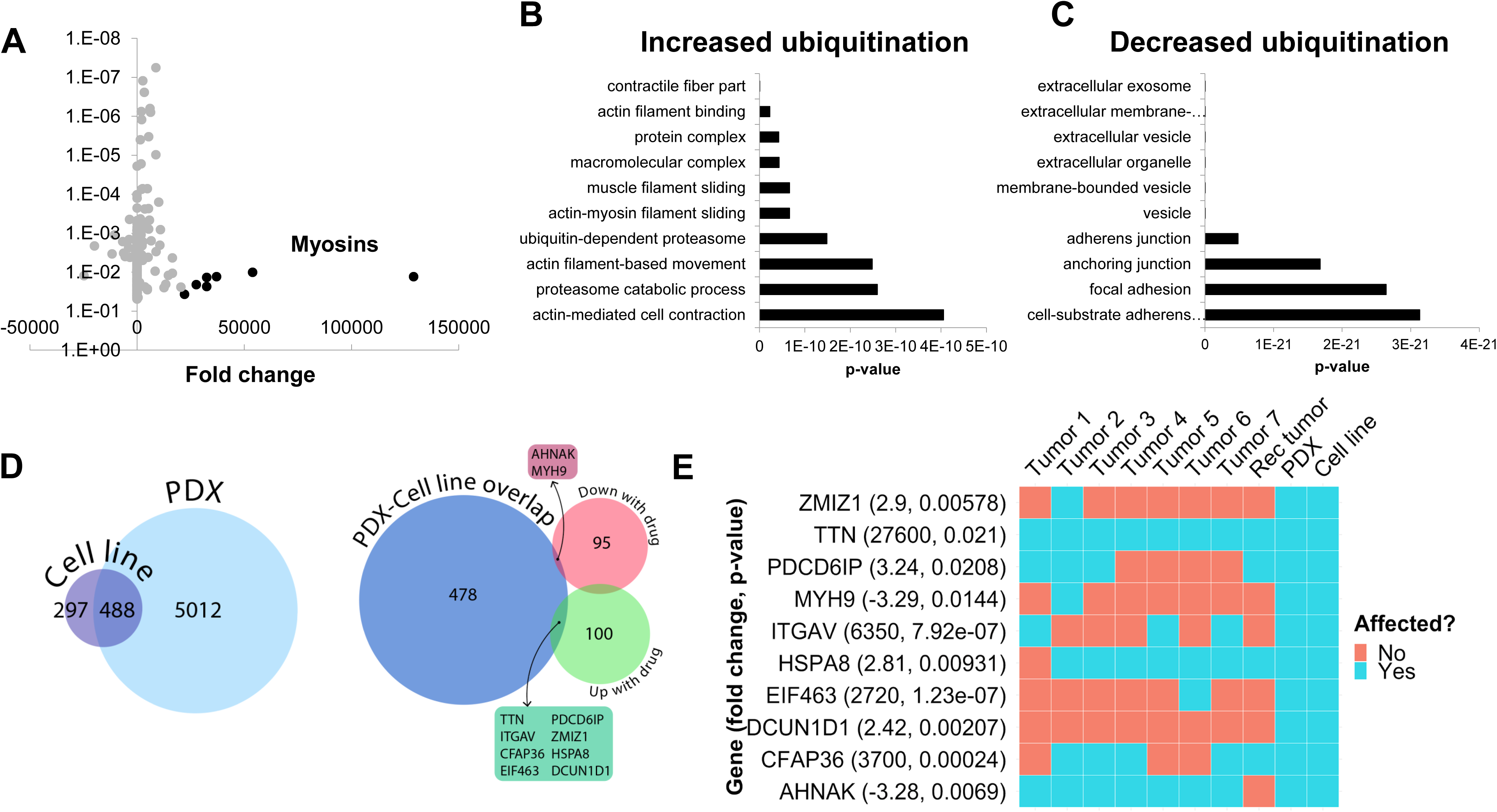
Ubiquitin proteomics of PDX tumors treated with bortezomib show differential ubiquitination in key pathways related to cytoskeletal dynamics and the proteasome. **A.** Mass spectroscopy proteomics of ubiquitin tagged proteins identified increased ubiquitination of 160 proteins and decreased ubiquitination of 130 proteins. Multiple myosins displayed increased ubiquitination in bortezomib-treated tumors. **B.** Pathway analysis of proteins with increased ubiquitination showed enrichment in pathways related to actin and proteasome subunits. **C.** Pathway analysis of proteins with decreased ubiquitination showed enrichment in pathways related to adherens junctions, focal adhesions, and extracellular vesicles. **D.** Genes affected by deleterious mutations in each sample were determined by analyzing the whole exome sequencing data with the Ensembl Variant Effect Predictor. Affected genes in the PDX were filtered by those in the cell line to eliminate potential contamination by mouse tissue (left). Comparison of this subset of genes with the proteins identified by proteomics analysis with increased or decreased ubiquitination in the PDXs treated with bortezomib identified an overlap of only ten affected proteins. **E.** The ten proteins identified in **D.** are shown and were affected in the tumors with high variability.

We next cross-referenced the proteomics analysis with our whole exome sequencing data to better understand the varied clinical response and rapid progression on bortezomib. We identified 10 proteins that contained identical somatic mutations across multiple samples predicted to alter protein function that were also differentially-ubiquitinated in the PDX and cell-line (**Figure 7D**). Interestingly, two of these 10 proteins are involved in pathways relevant to proteasome inhibition and HSPs, respectively (**Figure 7E**). Defective In Cullin Neddylation 1 Domain Containing 1 (DCUN1D1) is part of an E3 ubiquitin ligase complex for neddylation, and heat shock protein 70 kDa member 8 (HSPA8) is integral to the HSP70 pathway and cellular protein quality control systems (*33, 34*). Notably, the DCUN1D1 mutation was unique to the PDX and cell line (**Figure 7E**), suggesting the tumor from which this PDX was derived may have harbored unique genetics that could contribute to increased bortezomib sensitivity. Overall, the presence of somatic mutations affecting genes related to the proteasome and the heat shock protein pathway may explain the sensitivity to small molecule inhibitors targeting these pathways. The extensive heterogeneity in somatic mutations across multiple tumors and the patient-derived models may also help explain the rapid progression of the patient treated with the proteasome inhibitor, bortezomib.

## Discussion

### A comparative oncology approach enables rapid testing of a drug discovery pipeline in the clinic

Our canine leiomyosarcoma patient provided an invaluable opportunity to test, in real time, a personalized approach to cancer therapy. To do this, we generated patient-derived cancer models, both *in vitro* and *in vivo*, that helped identify novel therapeutic options, including proteasome inhibitors and HSP inhibitors. After identifying bortezomib as a potential drug for clinical application, we provided the preclinical data to the veterinary oncology team who initiated personalized therapy with bortezomib for local recurrence and metastatic disease. Though there was initial response to bortezomib in the setting of adjuvant palliative radiation therapy for the local recurrence, additional metastatic sites showed either stable disease or progression on bortezomib. While the outcome for this patient was only a slight delay in disease progression, the entire process of evaluating a personalized therapy – from presentation to death – was able to be carried out in approximately one year.

### The impact of genetic heterogeneity on treatment response

There are a number of possibilities to explain the rapid progression on bortezomib for this patient. One possible cause is the potential genetic drift that could be associated with generation and passage of the PDX and cell line. Indeed, recent studies have shown that PDXs are subject to mouse-specific selective pressures beyond a few passages (*9*). While we strive to keep our passage numbers low for this reason, it is possible that even the first implantation of a tumor into mice leads to selection of a specific sub-clone that has different properties from the original tumor. Interestingly, phylogenetic reconstruction of all seven tumors, a recurrent tumor and the PDX/cell line supports the grouping of the PDX/cell line with the recurrent tumor in a distinct clade. Consistent with this grouping, a recurrent tumor, like the PDX and cell line, had an initial response to bortezomib (**Figure 5**).

One additional possibility for the rapid clinical progression on bortezomib could be that there is not an established dosage or dosing schedule for treating canine cancer with bortezomib. Bortezomib has been used in veterinary medicine as a therapy for golden retriever muscular dystrophy and our therapeutic regimen was extrapolated from this (*35*). However, it is possible our dosing regimen was incorrect in the context of leiomyosarcoma treatment.

A third possibility is that the recurrent and metastatic lesions acquired unique mutations in key cellular pathways that conferred bortezomib resistance. Tumors are heterogeneous on the individual level and within the population, greatly contributing to the challenge of discovering novel universal drugs (*37–39*). Numerous studies across multiple cancer types have revealed significant genotypic variability even within a single tumor (*40, 41*) (*42–44*). This is the case for metastatic progression as well. For example, Wu et al. have shown that genetic signatures of metastatic lesions are similar to each other, but distinct from primary tumors, suggesting key genomic differences that could impact therapeutic response (*45*).

Driven by selective pressure from the tumor microenvironment, the inter-tumoral heterogeneity exhibited by these tumors could explain the difference between the *in vivo* response to bortezomib and the lack of response in the recurrent and metastatic lesions (*46, 47*). Consistent with this hypothesis, our analysis of whole exome sequencing data revealed substantial tumor heterogeneity across the multiple tumors from this patient, as well as between the group of samples including the recurrent tumor, PDX, and cell line.

It is possible that heterogeneity-mediated differences in response to therapy could be addressed with combination targeted therapy or with therapies that target multiple oncogenic pathways simultaneously (*39, 48, 49*). Multiple studies in mouse models of EGFR mutant lung cancer have shown the utility of combination therapies in overcoming treatment resistance (*50–52*). Our 2,100-compound drug screen identified multiple candidate drugs with both single cellular targets and those that target multiple pathways. In future iterations of this personalized pipeline, using combination therapy of top drugs identified from the drug screen could yield promising results.

### A multi-omics analysis identifies mechanisms of sensitivity and resistance to bortezomib

Using whole exome sequencing we were able to characterize the genomic differences between the tumor used for preclinical modeling and the recurrent tumors treated with bortezomib. In the context of multiple myeloma, for which bortezomib is a standard therapy, multiple cellular pathways have been associated with bortezomib resistance, including mutations in genes regulating the active site for bortezomib (*53–56*). Our proteomics analysis identified pathways related to actin-myosin filaments, HSPs, and the proteasome as downregulated by bortezomib (**Figure 7** and **Supplementary File 1**). The downregulation of skeletal myosins (MYH1, MYH2, MYH4) by bortezomib is not easily explained, since skeletal myosins are typically markers of rhabdomyosarcoma rather than leiomyosarcoma (*58*). However, inhibition of pathways related to HSPs and the proteasome further validates the target specificity and mechanism of action for bortezomib. Our integrated comparison of the ubiquitin proteomics data with the exome sequencing data identified 10 key genes that were both differentially ubiquitinated and mutated. Remarkably, two of these genes are members of the HSP and proteasome pathways. This integrated multi-omics analysis suggests that mutations within these two genes may explain, in part, the response to bortezomib. Likewise, the lack of mutation in these two genes within other tumors in this patient may also explain the differential response to bortezomib in different metastatic lesions of this patient.

## Conclusions

We have developed a translational drug discovery pipeline that integrates patient-derived models of cancer, drug screening, genomics, and proteomics to provide a comprehensive view of how to integrate translational preclinical research in the clinic. The unique biology of Teddy, with multiple, synchronous leiomyosarcoma tumors and an aggressive clinical course, enabled us to study the relationships between the molecular/genomic landscape and *in vitro, in vivo,* and clinical response to therapy. This provided both the patient and the clinician with unique information about tumor biology and response to novel therapeutics occurring in a very short period of time. This suggests that utilizing pet dogs with cancer to model personalized medicine approaches can facilitate rapid investigations of therapeutic successes and failures.

## Acknowledgments

JAS acknowledges support from Meg and Bill Lindenberger, the Paul and Shirley Friedland Fund, the Triangle Center for Evolutionary Medicine, and funds raised in memory of Muriel E. Rudershausen (riding4research.org). We would like to thank Teddy and his family for their participation in this study and the veterinary team at the University of Illinois at Urbana–Champaign for contributing to Teddy’s clinical care. We acknowledge Wayne Glover for contributing to *in vivo* PDX tumor propagation, the Duke Functional Genomics Shared Resource, the Duke Proteomics and Metabolomics Shared Resource, the Duke Sequencing and Genomics Shared Resource, and the Duke Genomic Analysis and Bioinformatics Core Facility.

## References

1. J. A. DiMasi, R. W. Hansen, H. G. Grabowski, The price of innovation: new estimates of drug development costs. J Health Econ 22, 151–185 (2003).

2. L. Mirabello, R. J. Troisi, S. A. Savage, Osteosarcoma incidence and survival rates from 1973 to 2004: data from the Surveillance, Epidemiology, and End Results Program. Cancer 115, 1531–1543 (2009).

3. T. A. C. S. m. a. e. c. team. (2018), vol. 2018.

4. D. F. Quail, J. A. Joyce, Microenvironmental regulation of tumor progression and metastasis. Nat Med 19, 1423–1437 (2013).

5. A. Blomme et al., Murine stroma adopts a human-like metabolic phenotype in the PDX model of colorectal cancer and liver metastases. Oncogene 37, 1237–1250 (2018).

6. J. J. Tentler et al., Patient-derived tumour xenografts as models for oncology drug development. Nature Reviews Clinical Oncology 9, 338–350 (2012).

7. Z. Chen et al., A murine lung cancer co-clinical trial identifies genetic modifiers of therapeutic response. Nature 483, 613–617 (2012).

8. I. Fichtner et al., Anticancer drug response and expression of molecular markers in early-passage xenotransplanted colon carcinomas. Eur J Cancer 40, 298–307 (2004).

9. M. Hidalgo et al., Patient-Derived Xenograft Models: An Emerging Platform for Translational Cancer Research. Cancer Discovery 4, 998 (2014).

10. M. Hidalgo et al., A pilot clinical study of treatment guided by personalized tumorgrafts in patients with advanced cancer. Mol Cancer Ther 10, 1311–1316 (2011).

11. J. Stebbing et al., Patient-derived xenografts for individualized care in advanced sarcoma. Cancer 120, 2006–2015 (2014).

12. N. Ehrhart, Soft-tissue sarcomas in dogs: a review. J Am Anim Hosp Assoc 41, 241–246 (2005).

13. C. A. Stiller et al., Descriptive epidemiology of sarcomas in Europe: report from the RARECARE project. Eur J Cancer 49, 684–695 (2013).

14. J. M. Dobson, S. Samuel, H. Milstein, K. Rogers, J. L. Wood, Canine neoplasia in the UK: estimates of incidence rates from a population of insured dogs. J Small Anim Pract 43, 240–246 (2002).

15. M. Milovancev et al., Comparative pathology of canine soft tissue sarcomas: possible models of human non-rhabdomyosarcoma soft tissue sarcomas. J Comp Pathol 152, 22–27 (2015).

16. J. M. Uronis et al., Histological and molecular evaluation of patient-derived colorectal cancer explants. PLoS One 7, e38422 (2012).

17. J. K. Cooper et al., Species identification in cell culture: a two-pronged molecular approach. In Vitro Cellular & Developmental Biology - Animal 43, 344–351 (2007).

18. Y. Shapovalov, D. Benavidez, D. Zuch, R. A. Eliseev, Proteasome inhibition with bortezomib suppresses growth and induces apoptosis in osteosarcoma. Int J Cancer 127, 67–76 (2010).

19. Y. Hu, D. Bobb, J. He, D. A. Hill, J. S. Dome, The HSP90 inhibitor alvespimycin enhances the potency of telomerase inhibition by imetelstat in human osteosarcoma. Cancer Biol Ther 16, 949–957 (2015).

20. B. Bioinformatics. (2018), vol. 2018.

21. M. Martin, Cutadapt Removes Adapter Sequences From High-Throughput Sequencing Reads. EMBnet.journal 17, (2011).

22. G. A. Van der Auwera et al., From FastQ data to high confidence variant calls: the Genome Analysis Toolkit best practices pipeline. Curr Protoc Bioinformatics 43, 11 10 11–33 (2013).

23. H. Li, R. Durbin, Fast and accurate short read alignment with Burrows-Wheeler transform. Bioinformatics 25, 1754–1760 (2009).

24. B. Institute. (Github), vol. 2018.

25. P. J. Kersey et al., Ensembl Genomes: an integrative resource for genome-scale data from non-vertebrate species. Nucleic Acids Res 40, D91–97 (2012).

26. A. McKenna et al., The Genome Analysis Toolkit: a MapReduce framework for analyzing next-generation DNA sequencing data. Genome Res 20, 1297–1303 (2010).

27. J. Felsenstein, Using the quantitative genetic threshold model for inferences between and within species. Philos Trans R Soc Lond B Biol Sci 360, 1427–1434 (2005).

28. E. Paradis, K. Schliep, ape 5.0: an environment for modern phylogenetics and evolutionary analyses in R. Bioinformatics, (2018).

29. P. Danecek et al., The variant call format and VCFtools. Bioinformatics 27, 2156–2158 (2011).

30. D. R. Zerbino et al., Ensembl 2018. Nucleic Acids Res 46, D754–D761 (2018).

31. T. Hulsen, J. de Vlieg, W. Alkema, BioVenn - a web application for the comparison and visualization of biological lists using area-proportional Venn diagrams. BMC Genomics 9, 488 (2008).

32. J. R. Conway, A. Lex, N. Gehlenborg, UpSetR: an R package for the visualization of intersecting sets and their properties. Bioinformatics 33, 2938–2940 (2017).

33. F. Wang et al., Blocking nuclear export of HSPA8 after heat shock stress severely alters cell survival. Sci Rep 8, 16820 (2018).

34. D. C. Scott et al., Blocking an N-terminal acetylation-dependent protein interaction inhibits an E3 ligase. Nat Chem Biol 13, 850–857 (2017).

35. K. P. Araujo et al., Bortezomib (PS-341) treatment decreases inflammation and partially rescues the expression of the dystrophin-glycoprotein complex in GRMD dogs. PLoS One 8, e61367 (2013).

36. R. Fisher, L. Pusztai, C. Swanton, Cancer heterogeneity: implications for targeted therapeutics. Br J Cancer 108, 479–485 (2013).

37. N. McGranahan, C. Swanton, Biological and therapeutic impact of intratumor heterogeneity in cancer evolution. Cancer Cell 27, 15–26 (2015).

38. J. Liu, H. Dang, X. W. Wang, The significance of intertumor and intratumor heterogeneity in liver cancer. Exp Mol Med 50, e416 (2018).

39. I. Dagogo-Jack, A. T. Shaw, Tumour heterogeneity and resistance to cancer therapies. Nat Rev Clin Oncol 15, 81–94 (2018).

40. B. A. Walker et al., Intraclonal heterogeneity and distinct molecular mechanisms characterize the development of t(4;14) and t(11;14) myeloma. Blood 120, 1077–1086 (2012).

41. M. Gerlinger et al., Intratumor heterogeneity and branched evolution revealed by multiregion sequencing. N Engl J Med 366, 883–892 (2012).

42. A. Sottoriva et al., Intratumor heterogeneity in human glioblastoma reflects cancer evolutionary dynamics. Proc Natl Acad Sci U S A 110, 4009–4014 (2013).

43. S. Bea et al., Landscape of somatic mutations and clonal evolution in mantle cell lymphoma. Proc Natl Acad Sci U S A 110, 18250–18255 (2013).

44. A. Kogita et al., Inter-and intra-tumor profiling of multi-regional colon cancer and metastasis. Biochem Biophys Res Commun 458, 52–56 (2015).

45. X. Wu et al., Clonal selection drives genetic divergence of metastatic medulloblastoma. Nature 482, 529–533 (2012).

46. M. R. Junttila, F. J. de Sauvage, Influence of tumour micro-environment heterogeneity on therapeutic response. Nature 501, 346–354 (2013).

47. M. S. Lawrence et al., Mutational heterogeneity in cancer and the search for new cancer-associated genes. Nature 499, 214–218 (2013).

48. A. A. Alizadeh et al., Toward understanding and exploiting tumor heterogeneity. Nat Med 21, 846–853 (2015).

49. Z. Liao et al., The Anthelmintic Drug Niclosamide Inhibits the Proliferative Activity of Human Osteosarcoma Cells by Targeting Multiple Signal Pathways. Curr Cancer Drug Targets 15, 726–738 (2015).

50. V. Pirazzoli et al., Afatinib plus Cetuximab Delays Resistance Compared to Single-Agent Erlotinib or Afatinib in Mouse Models of TKI-Naive EGFR L858R-Induced Lung Adenocarcinoma. Clin Cancer Res 22, 426–435 (2016).

51. Y. Y. Janjigian et al., Dual Inhibition of EGFR with Afatinib and Cetuximab in Kinase Inhibitor-Resistant EGFR-Mutant Lung Cancer with and without T790M Mutations. Cancer Discovery 4, 1036–1045 (2014).

52. E. M. Tricker et al., Combined EGFR/MEK Inhibition Prevents the Emergence of Resistance in EGFR-Mutant Lung Cancer. Cancer Discov 5, 960–971 (2015).

53. R. Oerlemans et al., Molecular basis of bortezomib resistance: proteasome subunit beta5 (PSMB5) gene mutation and overexpression of PSMB5 protein. Blood 112, 2489–2499 (2008).

54. D. Chauhan et al., Blockade of Hsp27 overcomes Bortezomib/proteasome inhibitor PS-341 resistance in lymphoma cells. Cancer Res 63, 6174–6177 (2003).

55. D. J. Kuhn et al., Targeting the insulin-like growth factor-1 receptor to overcome bortezomib resistance in preclinical models of multiple myeloma. Blood 120, 3260–3270 (2012).

56. W. Que, J. Chen, M. Chuang, D. Jiang, Knockdown of c-Met enhances sensitivity to bortezomib in human multiple myeloma U266 cells via inhibiting Akt/mTOR activity. APMIS 120, 195–203 (2012).

57. T. Saku, N. Tsuda, M. Anami, H. Okabe, Smooth and skeletal muscle myosins in spindle cell tumors of soft tissue. An immunohistochemical study. Acta Pathol Jpn 35, 125–136 (1985).

