## Supplementary Figure Legends for "From the clinic to the bench and back again in one dog year: identifying new treatments for sarcoma using a cross-species personalized medicine pipeline"

**Supplementary Figure 1.** **Radiographic and physical exam findings demonstrating the initial presentation of local (right pelvic limb) and metastatic (nasal lesion, tracheobronchial lymph nodes) sites in comparison to images obtained after bortezomib treatment.** Overall the lesions appear larger after one month of systemic bortezomib therapy with the nasal lesion infiltrating the maxilla and overall increase in size and heterogeneity of the pelvic and lymph node lesions.

**Supplementary Figure 2. Phylogenetic reconstruction of Teddy’s tumors and patient-derived models using A. maximum likelihood, B., maximum parsimony, and C. distance methods.** Maximum likelihood and maximum parsimony algorithms group the PDX and cell line with the recurrent tumor with high (>90%) bootstrap support. The distance tree places the PDX with tumor 1. Clades with >90% bootstrap support are highlighted in teal.
