## Supplementary Figure 1 for "From the clinic to the bench and back again in one dog year: identifying new treatments for sarcoma using a cross-species personalized medicine pipeline"

### Before Bortezomib Treatment

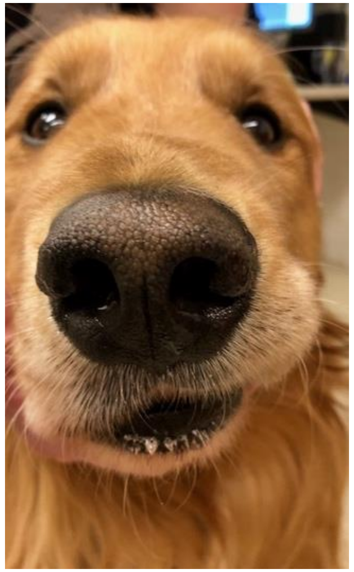

Nasal Lesion

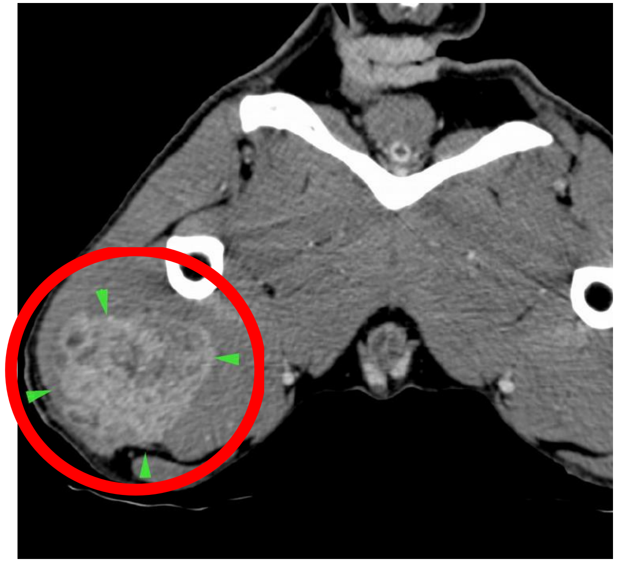

Recurrence at the right pelvic limb

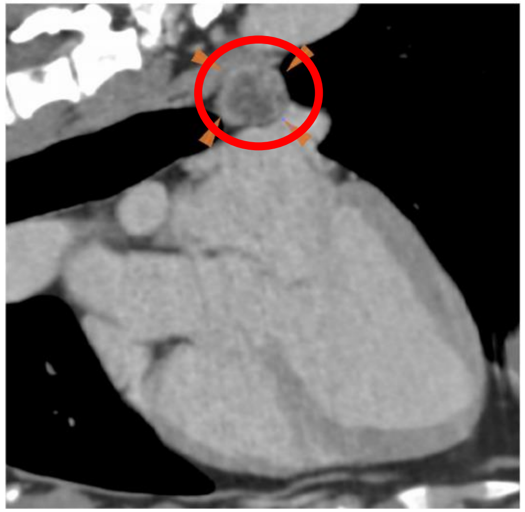

Tracheobronchial Lymph Node Metastasis

### After Bortezomib Treatment

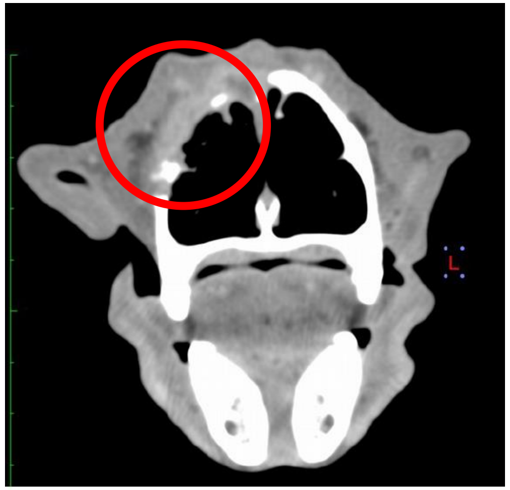

Nasal Lesion

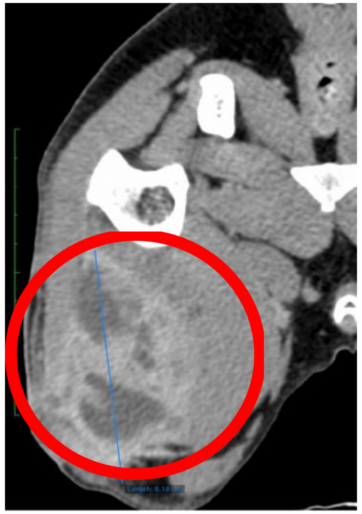

Recurrence at the right pelvic limb

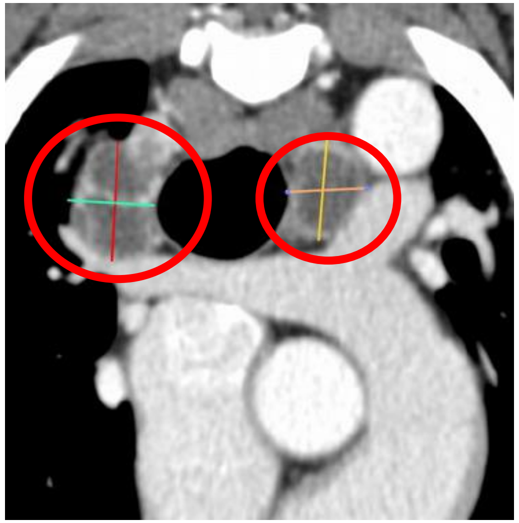

Tracheobronchial Lymph Node Metastasis
