## Supplementary figures and images for "From the clinic to the bench and back again in one dog year: identifying new treatments for sarcoma using a cross-species personalized medicine pipeline"

### Supplementary Figure 2

#### Supplementary Figure 2

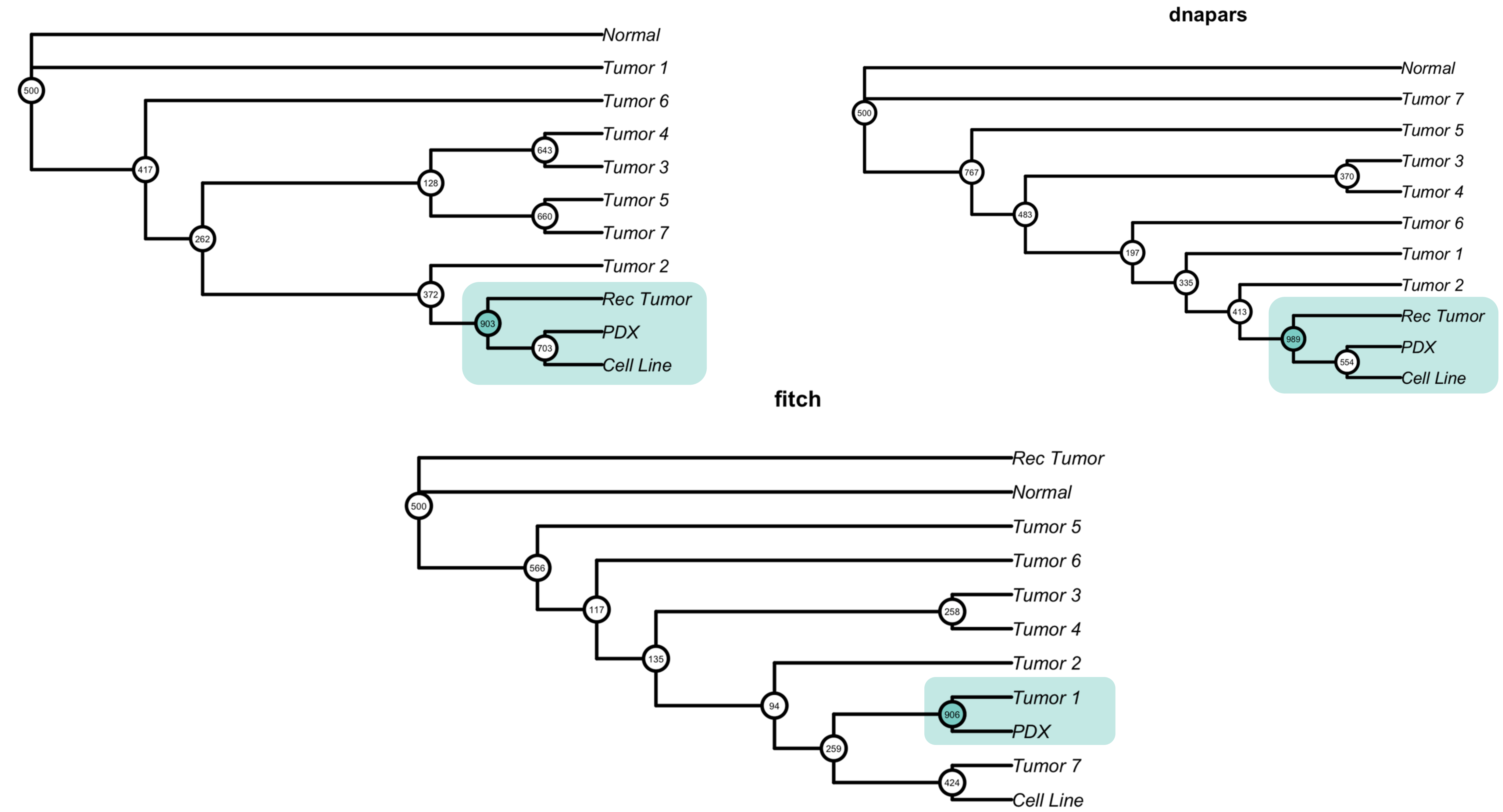
